# Histopathological Evaluation of Pulmonary Arterial Remodeling in COVID-19

**DOI:** 10.1101/2024.12.12.628253

**Authors:** Sergiy G. Gychka, Sofiia I. Nikolaienko, Nataliia V. Shults, Volodymyr M. Vasylyk, Bohdan O. Pasichnyk, Iryna V. Kagan, Yulia V. Dibrova, Muin Tuffaha, Yuichiro J. Suzuki

## Abstract

A positive-sense single-stranded RNA virus, severe acute respiratory syndrome coronavirus 2 (SARS-CoV-2), caused the coronavirus disease 2019 (COVID-19) pandemic that devastated the world. While this is a respiratory virus, one feature of the SARS-CoV-2 infection was recognized to cause pathogenesis of other organs. Because the membrane fusion protein of SARS-CoV-2, the spike protein, binds to its major host cell receptor angiotensin-converting enzyme 2 (ACE2) that regulates a critical mediator of cardiovascular diseases, angiotensin II, COVID-19 is largely associated with vascular pathologies. In fact, we have previous reported that postmortem lung tissues collected from patients who died of COVID-19 exhibited thickened pulmonary vascular walls and reduced vascular lumen. The present study extended these findings by further characterizing the pulmonary vasculature of COVID-19 patients using larger sample sizes and providing mechanistic information through histological observations. The examination of 56 autopsy lung samples showed thickened vascular walls of small pulmonary arteries after 14 days of disease compared to H1N1 influenza patients who died before COVID- 19 pandemic started. Pulmonary vascular remodeling in COVID-19 patients was associated with hypertrophy of the smooth muscle layer, perivascular fibrosis, edema and lymphostasis, inflammatory infiltration, perivascular hemosiderosis and neoangiogenesis. We found a correlation between the duration of hospital stay and the thickness of the muscular layer of pulmonary arterial walls. These results further confirm that COVID-19 affects the pulmonary vasculature and warrants an evaluation of patients that survived COVID-19 for possible future development of pulmonary arterial hypertension.

## Introduction

Coronaviruses are positive-sense single-stranded RNA viruses that often cause the mild common cold [1,2]. However, some coronaviruses can be deadly. The world is still affected by the coronavirus disease 2019 (COVID-19) caused by severe acute respiratory syndrome coronavirus 2 (SARS-CoV-2) [3–5]. Nearly a billion people have become infected with SARS-CoV-2 worldwide, with millions of deaths, and COVID-19 has caused serious health, economic, and sociological problems. SARS-CoV-2 uses angiotensin-converting enzyme 2 (ACE2), a physiological regulator of blood pressure that converts angiotensin II (Ang II) to angiotensin 1-7 (Ang 1-7), as a receptor to enter the host cells for infection [6–8]. Lung cells are the primary targets of SARS-CoV-2, resulting in severe pneumonia and acute respiratory distress syndrome [9,10]. Certain populations of infected individuals are more susceptible to being severely affected by COVID-19. Elderly patients are particularly susceptible to developing severe and fatal conditions [4,11,12], suggesting that this virus also affects organs other than the respiratory system.

We have previously reported that postmortem lung tissues collected from 10 patients who died of COVID-19 in Ukraine during the period of March to July 2020 exhibited thickened pulmonary vascular walls compared to the archival materials of lung tissues from 17 patients who died of influenza A (H1N1) during the epidemic in November and December of 2009 [13]. In addition to these observations in postmortem tissues, we also found that the placental arteries of healthy women who gave birth to live full-term newborns who tested positive for COVID-19 during pregnancy without developing serious symptoms, exhibited severe vascular wall thickening and the occlusion of the vascular lumen [14]. Thus, the vascular system is sensitive to being affected by COVID-19 even for those who are asymptomatic or only mildly symptomatic, and thus we need to consider the possibility of vascular problems post-pandemic. We previously communicated our concern that COVID-19 patients may become predisposed to pulmonary arterial hypertension with the scientific community [15]. Thus, further investigation into the effects of COVID-19 on the pulmonary vessels are needed.

In the present study, using autopsy lung samples from patients who died of COVID-19 as well as H1N1 influenza in Ukraine, we performed detailed histopathological characterization of the effects of COVID-19 on pulmonary vessels.

## Materials and Methods

### Patient Samples

Fifty-six postmortem autopsy lung samples from patients who died of COVID-19 with severe fatal course of disease were obtained in Ukraine during the period of March 2020 to January 2023. All cases were divided into three groups in accordance with the duration of the disease: Group 1 – death on or before the 8^th^ day of disease (18 cases); Group 2 – death between the 9^th^ and 13^th^ day of disease (20 cases); Group 3 – death after the 14^th^ day of disease (18 cases); and Group 4 (control group) – patients who died of H1N1 influenza in 2009 between the 7^th^ and 14^th^ day of disease (17 cases). The onset of the disease was established by a clinical diagnosis of COVID-19 or H1N1 influenza. These viral infections were confirmed by PCR tests. Autopsy cases used in this study were randomly selected, and the time of autopsy did not exceed 12 hours after death. In this study, the average age of COVID-19 and H1N1 patients at the time of death was 62.2 years of age. 49% of COVID-19 patients were women. These studies were approved by the regional committee for medical research ethics in Kyiv, Ukraine and performed under the Helsinki Declaration of 1975 revised in 2013 or comparable ethical standards.

### Histological Examinations

Autopsy sample material approximately 10 mm thick were fixed overnight in 10% buffered formalin at room temperature. Fixed tissues were embedded in paraffin. From paraffin blocks, 5 µm thick sections were made using a microtome. The sections were subjected to hematoxylin and eosin, Masson’s trichrome, and Verhoeff Van Gieson (EVG) stainings. Immunohistochemistry was also performed using anti-AXL (Abcam Limited, Cambridge, UK), anti-spike protein (MyBioSource, Inc., San Diego, CA, USA), anti-α-smooth muscle actin (Thermo Fisher Scientific, Waltham, MA, USA), anti-CD34 (VITRO S.A., Alcalá de Guadaíra, Spain), and anti-CD105 (Abcam) antibodies. Histological specimens were examined using an Axiolab 5 microscope with an Axiocam 305 color digital camera, and the Labscope v. 2.7 software. Morphometric investigations included the assessment of pulmonary arterial wall thickness, the ratio of internal pulmonary arterial area to external pulmonary arterial area, expressed both in hundredth (area index) and percent (lumen area %). Six (3 small-sized and 3 medium-sized) vessels were analyzed for each patient.

### Statistical Analysis

GraphPad Prism version 10.3.1 program (GraphPad Soft-ware, La Jolla, CA) was used for statistical analysis of the data. The results were statistically analyzed using one-way ANOVA followed by a Tukey’s test for multiple comparisons. A p value less than or equal to 0.05 was considered significant. Results are expressed as the mean ± SEM of n experiments. Significance was tested using two-tailed Mann–Whitney test and Kruskal-Wallis test (non-parametric one-way ANOVA) for multiple groups comparison.

## Results

Histological and immunohistochemical examination of autopsy materials of patients who died of COVID-19 with a severe fatal course of the disease demonstrated signs of SARS-CoV- 2-mediated damage to the lung structures, in particular the pulmonary arteries. Expression of ACE2 was detected in the endothelial cells, in particular in vasa vasorum (Fig. 1a). In the pulmonary arterial walls, the expression of AXL, that has been shown to be an alternative cell entry receptor for SARS-CoV-2 [16], was detected. The expression of AXL was observed, not only in alveolocytes and alveolar macrophages, but also in smooth muscle cells of the pulmonary arterial walls (Fig. 1b), suggesting the possible direct invasion of SARS-CoV-2 into smooth muscle cells. Indeed, while SARS-CoV-2 spike protein expression was largely observed in endothelial cells (Fig. 1c), in some cases, the spike protein was also detected in smooth muscle cells of pulmonary arterial walls (Fig.1d). Thus, in severe course of COVID-19, SARS-CoV-2 and its proteins are present in the structures of the pulmonary arterial walls.

**Fig. 1.**
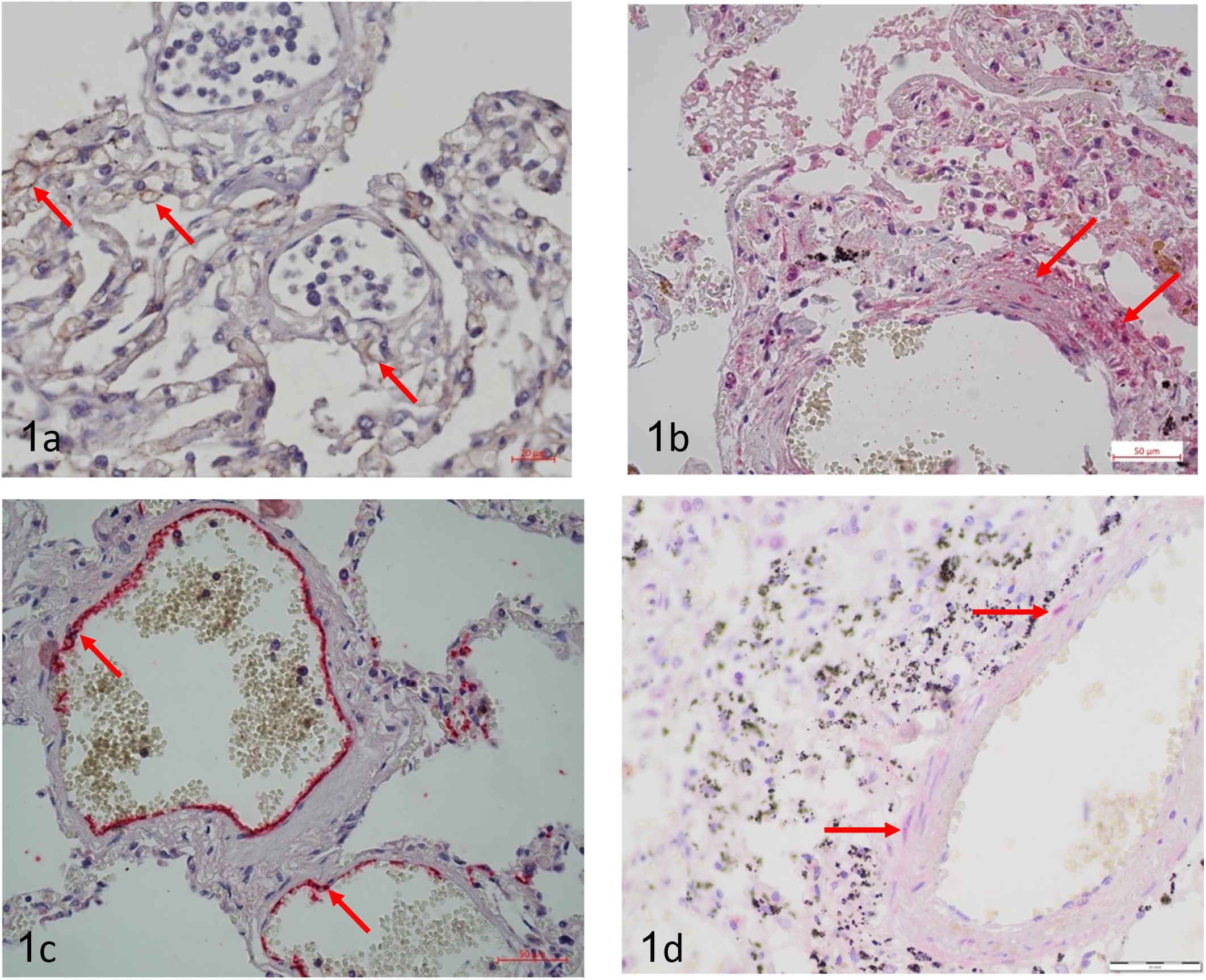
The expression of SARS-CoV-2 entry receptors and spike protein in pulmonary vessels. a – Expression of ACE2 as an receptor of SARS-CoV-2 entry in the endothelial cells of capillaries and vasa vasorum (red arrows). IHC with anti-ACE2. Scale bar – 20 μm. b – Expression of AXL as an alternative receptor of SARS-CoV-2 entry in smooth muscle cells (SMCs) of the pulmonary artery wall (red arrows). IHC with anti-AXL. Scale bar – 50 μm. c – Spike protein expression in the endothelial cells of the pulmonary arteries (red arrows). IHC with anti-Spike. Scale bar – 50 μm. d - Spike protein expression in SMCs of the pulmonary artery wall (red arrows). IHC with anti-Spike. Scale bar – 50 μm.

We have previously reported that the main histological manifestations of COVID-19, in addition to respiratory distress syndrome and pneumonia, were thickening of the pulmonary arteries walls [13]. This new study with a large number of patient samples confirmed earlier findings and determined that the severity of pulmonary arteries thickening depends on the duration of the disease. Marked remodeling of the pulmonary arteries was detected in patients who died of COVID-19 at least 2 weeks after the onset of the disease (Fig. 2). One of the morphological features of the pulmonary arteries was the thickening (hypertrophy) of the smooth muscle layer of their walls. Immunohistochemical analysis of the pulmonary arteries of patients who died of COVID-19 indicated that the thickening is largely due to the hypertrophy of the smooth muscle layer in small pulmonary arteries (Fig. 3a, 3b, 3c, 3d). We performed morphometric analysis to assess the differences between the four groups in two indicators: the thickness of the muscular layer of arteries measuring 50-100 μm (Fig. 3e, 3g) and 100-200 μm (Fig. 3f, 3h), and the lumen area and area index of both small-sized arteries (50-100 μm; Fig. 3i, 3k) and medium-sized arteries (100-200 μm; Fig. 3j, 3l).

**Fig. 2.**
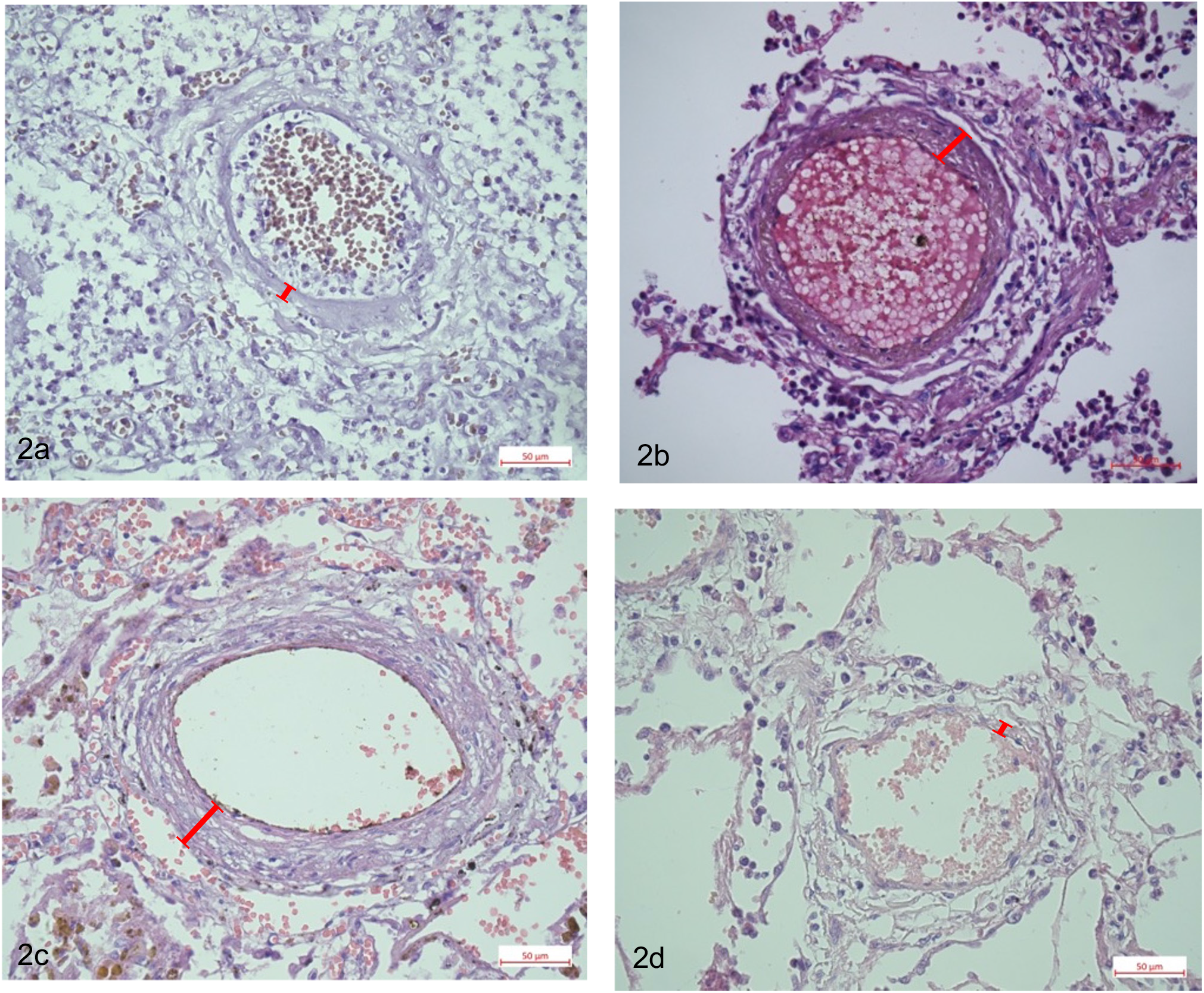
Thickening of pulmonary arteries walls in COVID-19 in comparison with H1N1 influenza. a – mild thickening of the medium-sized pulmonary artery wall in Group 1 – COVID- 19 till 8th day of disease. H&E staining. Scale bar – 50 μm. b – moderate thickening of the medium-sized pulmonary artery wall in Group 2 – COVID-19 between 9th and 13th day of disease. H&E staining. Scale bar – 50 μm. c – pronounced thickening of the medium-sized pulmonary artery wall in Group 3 – COVID-19 after 14th day of disease. H&E staining. Scale bar – 50 μm. d – no signs of thickening of the medium-sized pulmonary artery in Control Group (H1N1 influenza). H&E staining. Scale bar – 50 μm.

**Fig. 3.**
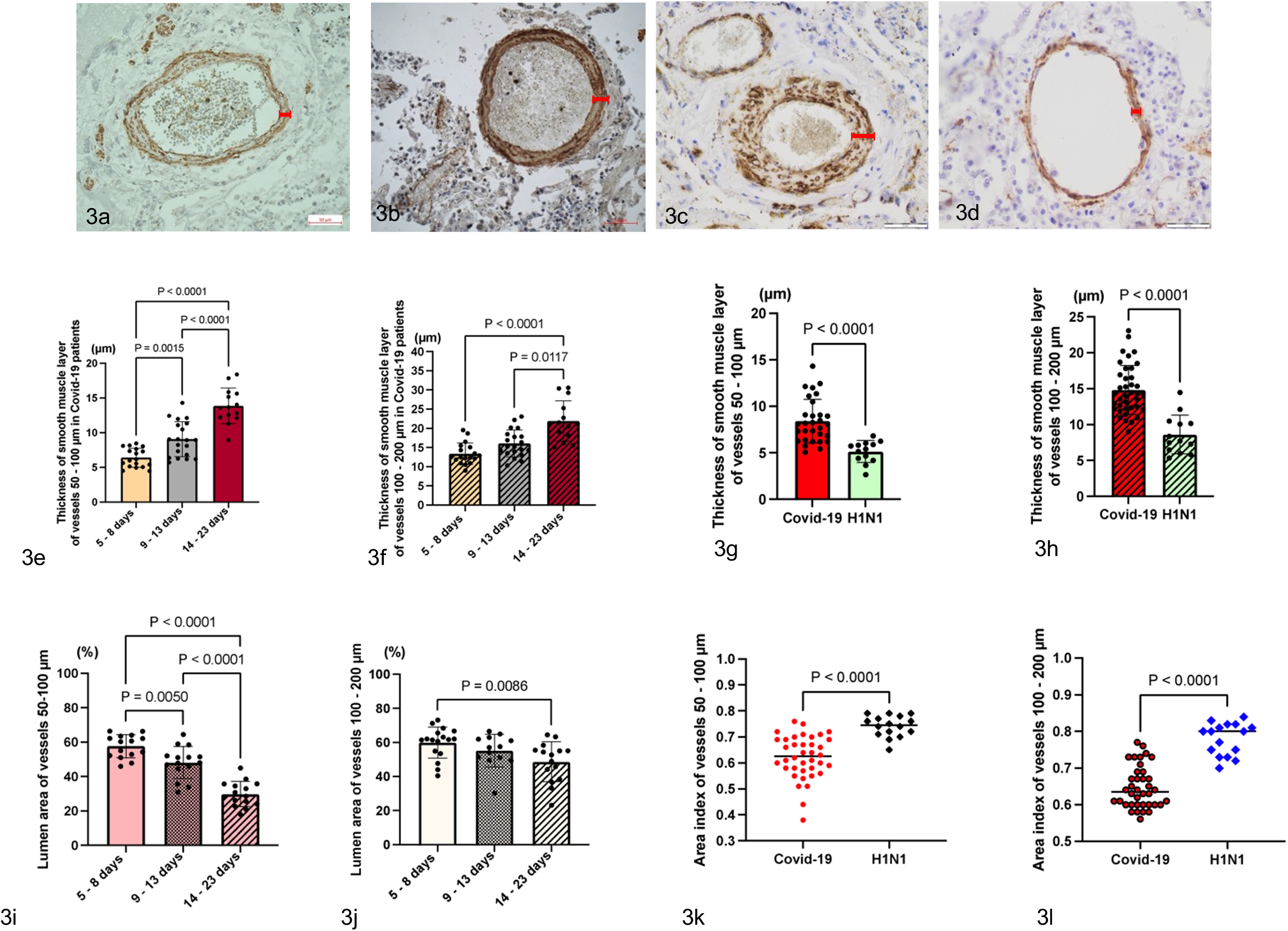
Hypertrophy of smooth muscle layer as a mechanism of thickening of the pulmonary arteries walls in COVID-19. a – mild hypertrophy of smooth muscle layer of the pulmonary artery wall in Group 1 – COVID-19 till 8th day of disease. IHC with sm-Actin. Scale bar – 50 μm. b – moderate hypertrophy of smooth muscle layer of the pulmonary artery wall in Group 2 – COVID-19 between 9th and 13th day of disease. IHC with sm-Actin. Scale bar – 50 μm. c – pronounced hypertrophy of smooth muscle layer of the pulmonary artery wall in Group 3 – COVID-19 after 14th day of disease. IHC with sm-Actin. Scale bar – 50 μm. d – no signs of hypertrophy of smooth muscle layer of the pulmonary artery wall in Control Group (H1N1 influenza). IHC with sm-Actin. Scale bar – 50 μm. Red arrows show thickness of muscular layer. e – the statistically significant difference was detected in the distribution of smooth muscle layer thickness of the small arteries between COVID-19 groups. Average for groups: Group 1=6,41 (StDev=1,37; CV=21,43%), Group 2=9,07 (StDev=2,52; CV=27,75%), Group 3=13,49 (StDev=2,52; CV=18,68%). f – the significant difference was detected in the distribution of muscular layer thickness of the medium-sized arteries between COVID-19 group. Average for groups: Group 1=13,42 (StDev=2,78; CV=20,72%), Group 2=16,08 (StDev=3,51; CV=21,81%), Group 3=23,49 (StDev=5,83; CV=24,84%). g - the significant difference was detected in the distribution of muscular layer thickness of the small arteries between all cases of COVID-19 group and H1N1 influenza control group. Average for groups: Group COVID-19 = 8.41 (StDev=2.34; CV=27.84%), H1N1 group= 5,13 (StDev=1,17; CV=22,76%). h - the significant difference was detected in the distribution of muscular layer thickness of the medium- sized arteries between all cases of COVID-19 group and H1N1 influenza control group. Average for groups: Group COVID-19 = 14.83 (StDev=3.46; CV=23.32%), H1N1 group=8,62 (StDev=2,72; CV=31,51%). i –the significant difference was detected in the distribution of lumen area (%) of the small arteries between COVID-19 groups. Average for groups: Group 1=55,38% (St.dev=0,09; CV=16,46%), Group 2=48,15% (St.dev=0,09; CV=19,33%), Group 3=32,58% (St.dev=0,1; CV=31,46%). j – the significant difference was detected in the distribution of lumen area (%) of the medium-sized arteries between COVID-19 groups. Average for groups: Group 1=59,96% (St.dev=0,09; CV=15,17%), Group 2=58,11% (St.dev=0,12; CV=19,96%), Group 3=50,78% (St.dev=0,12; CV=24,43%). k - the significant difference was detected in the distribution of area index of the small arteries between all cases of COVID-19 group and H1N1 influenza control group. Average for groups: Group COVID-19 = 0.62 (StDev=0.08; CV=13.34%), H1N1 group=0,74 (St.dev=0,04; CV=5,48%). l - the significant difference was detected in the distribution of area index of the medium-sized arteries between all cases of COVID-19 group and H1N1 influenza control group. Average for groups: Group COVID-19 = 0.65 (StDev=0.06; CV=8.82%), H1N1 group=0,78 (St.dev=0,04; CV=5,66%).

Statistically, the significant difference was detected in the distribution of lumen area (%) of the small and medium-sized arteries between COVID-19 groups (Fig. 3i, 3j). The percentage of the lumen area of the pulmonary arteries correlates with the duration of the disease. The smallest lumen area of both small and medium-sized vessels is observed in Group 3. For small vessels, statistical significance (p) between Group 1 and Group 2 is 0.0050, between Group 2 and Group 3 p < 0.0001, and between Group 1 and Group 3 p < 0.0001. Among medium-sized vessels, there was a significant difference between Group 1 and Group 3 (p = 0.0086). The highest wall thickness and the smallest lumen area index were observed in the group after 14th day.

Graphs 3k and 3l demonstrate a significant difference in the distribution of the area index of small and medium-sized vessels among all patients who died of COVID-19, compared with patients who died of H1N1 influenza (p < 0.0001).

COVID-19 may promote the development of perivascular fibrosis. The duration of the disease affects the severity of perivascular fibrosis through the inflammation caused by production of inflammatory cytokines and chemokines by immune cells and, as a result, the activation of fibroblasts that synthesize the collagen. COVID-19 was associated with the development of perivascular fibrosis in the pulmonary arteries of different grades – low, medium, and high – which depend on the duration of the disease (Fig. 4).

**Fig. 4.**
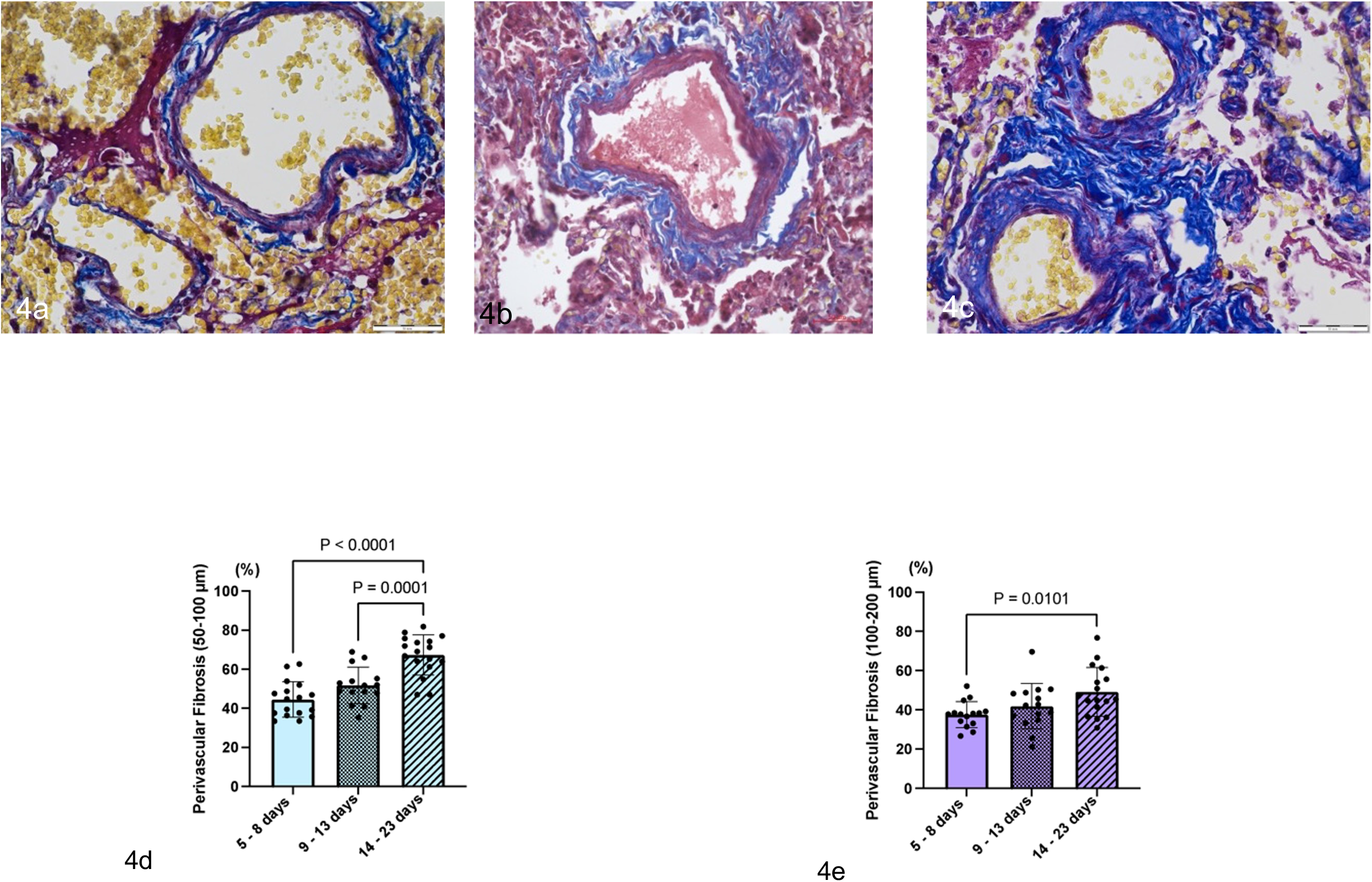
Perivascular fibrosis as a mechanism of pulmonary arteries thickening in COVID-19. a - low grade perivascular fibrosis in Group 1 - COVID-19 till 8^th^ day of disease. Masson’s Trichrome staining. Scale bar - 50 μm. b - medium grade perivascular fibrosis in Group 2 – COVID-19 between 9^th^ and 13^th^ day of disease. Masson’s Trichrome staining. Scale bar - 50 μm. c – high grade perivascular fibrosis in Group 3 - COVID-19 after 14^th^ day of disease. Masson’s Trichrome staining. Scale bar - 50 μm. d – the statistically significant difference was detected in the distribution of area of perivascular fibrosis of the small arteries between COVID- 19 groups. Average for groups: Group 1= 44,62% (StDev=0,09; CV=20,43%), Group 2=51,85% (StDev=0,09; CV=17,96%), Group 3=67,42% (StDev=0,10; CV=15,20%). e – the significant difference was detected in the distribution of perivascular fibrosis of the medium-sized arteries between COVID-19 group. Average for groups: Group 1=40,04% (StDev=9,09%; CV=22,72%), Group 2=41,89% (StDev=11,60%; CV=27,69%), Group 3=49,22% (StDev=12,40%; CV=25,20%).

In COVID-19, the lungs develop swelling of the parenchyma, including the pulmonary arteries walls, which causes decompensation of the lymph outflow. Lymphostasis is one of the factors of the arterial wall thickening due to the accumulation of proteins, detritus, and inflammatory cells in the perivascular space. As a result, fibroblasts are activated with the subsequent formation of connective tissue and the development of perivascular fibrosis which causes arterial wall thickening (Fig. 5a, 5b).

**Fig. 5.**
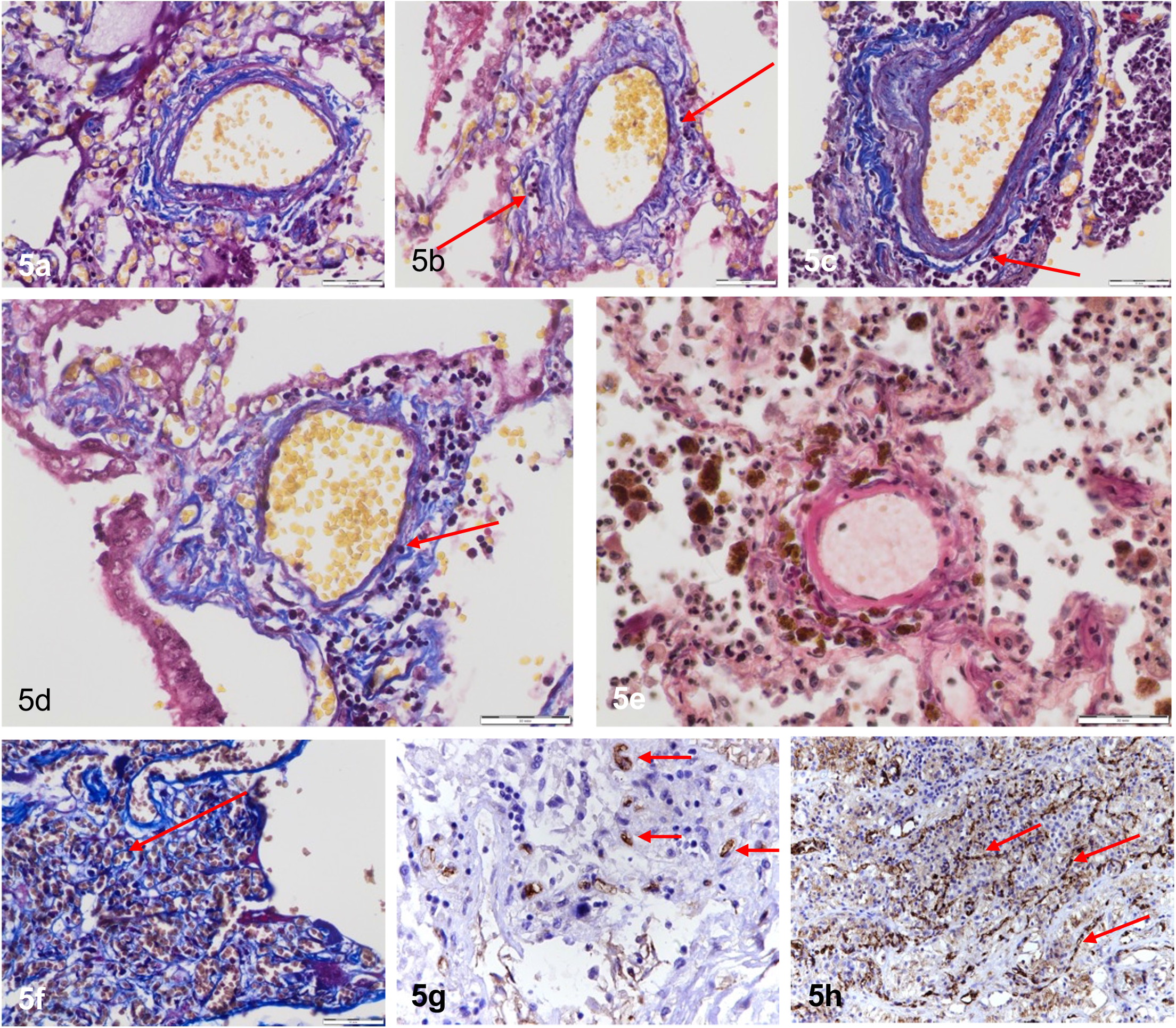
Other factors contributing to the pulmonary arterial thickening in COVID-19. a – swelling of pulmonary arterial wall. Masson’s Trichrome staining. Scale bar - 50 μm. b – lymphostasis. Red arrows show expanded lymphatic vessels in pulmonary arterial wall. Masson’s Trichrome staining. Scale bar - 50 μm. c – initial stage of vasculitis development. Red arrow shows inflammatory cells in perivascular area. Masson’s Trichrome staining. Scale bar - 50 μm. d – migration of inflammatory cells into the pulmonary arterial wall. Red arrow shows migration of inflammatory cells into the pulmonary arterial wall. Masson’s Trichrome staining. Scale bar - 50 μm. e - perivascular hemosiderosis. EVG staining. Scale bar - 50 μm. f – red arrows show newly-formed vessels. Masson’s Trichrome staining. Scale bar - 50 μm. g – red arrows show endothelium of newly-formed vessels detected by IHC with CD-105. x400. h – red arrows show newly-formed vessels detected by IHC with CD-31. x200.

Another factor for arterial wall thickening was perivascular inflammatory infiltration and vasculitis, which were detected in 27.8% of all COVID-19 cases (Fig. 5c, 5d). Statistical processing of the data showed no difference between the COVID-19 groups, indicating that the duration of the disease does not play a significant role in the development of vasculitis. Inflammatory infiltration of perivascular spaces is primarily due to activation of cell-mediated and humoral immunity and is manifested by type 3 hypersensitivity reactions [17]. Perivascular fibrosis and fibrosis of pulmonary arteries walls develop as a result of inflammatory damage of arterial wall tissues.

In some cases of COVID-19, we observed pronounced hemosiderosis in the perivascular areas around the pulmonary arteries (Fig. 5e). Hemosiderosis occurs as a result of prolonged and widespread hemorrhages in the alveoli. Alveolar macrophages phagocyte red blood cells and produce a significant amount of hemosiderin, becoming so-called siderophages. These siderophages are transported by lymph flow to the perivascular spaces of the pulmonary arteries. Alveolar macrophages with intracytoplasmic ferritin are known to cause oxidative stress, which also can lead to pulmonary fibrosis and perivascular fibrosis [18].

Neoangiogenesis may also be a factor that contributes to the thickening of pulmonary arteries. In COVID-19, we observed activation of neoangiogenesis in the perivascular areas around the pulmonary arteries. We found high expression of endoglin (CD105), which is a part of the TGFβ receptor complex of the membrane protein located on the surface of new-formed endothelial cells. Neoangiogenesis is associated with perivascular fibrosis, also leading to thickening of the arterial walls (Fig. 5f, 5g, 5h).

Pulmonary arterial thrombosis was observed in 42.6% of all cases without a significant difference between the study groups. Vasa vasorum thrombosis was detected in 8.2% of all cases (Fig. 6). Furthermore, SARS-CoV-2-induced vasa vasorum thrombosis led to the development of hypoxic stress and damage of the pulmonary arteries walls which cause their remodeling, fibrosis, and thickening.

**Fig. 6.**
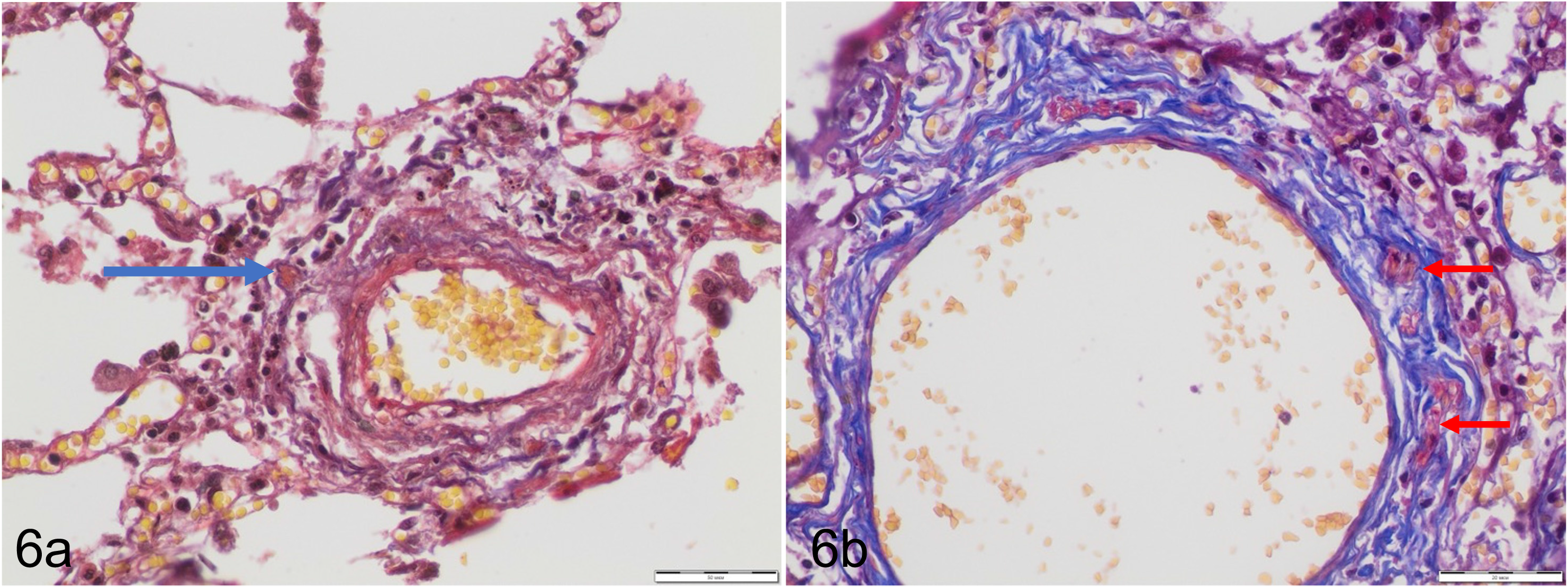
Vasa vasorum thrombosis leads to hypoxia of pulmonary arterial wall, which subsequently causes pulmonary arterial wall remodelling in COVID-19. a – blue arrow shows thrombus in the vasa vasorum lumen. EVG staining. Scale bar - 50 μm. b – red arrows show thrombi in the vasa vasorum lumens. Masson’s Trichrome staining. Scale bar - 20 μm.

Finally, we found a correlation between the duration of hospital stay and the thickness of the muscular layer of the pulmonary arteries walls (Fig. 7). Figures 7a and 7b demonstrate the thickness of the smooth muscle layer of the vessels over a 14-day hospitalization period. The thickness of the smooth muscle layer significantly increased in COVID-19 patients by day 14 of the disease compared to H1N1 patients. The AUC analysis (Figures 7c and 7d) indicates that COVID-19 patients experience a greater cumulative increase in smooth muscle layer thickness over the 14-day hospitalization period compared to H1N1 patients, with this difference being statistically significant.

**Fig. 7.**
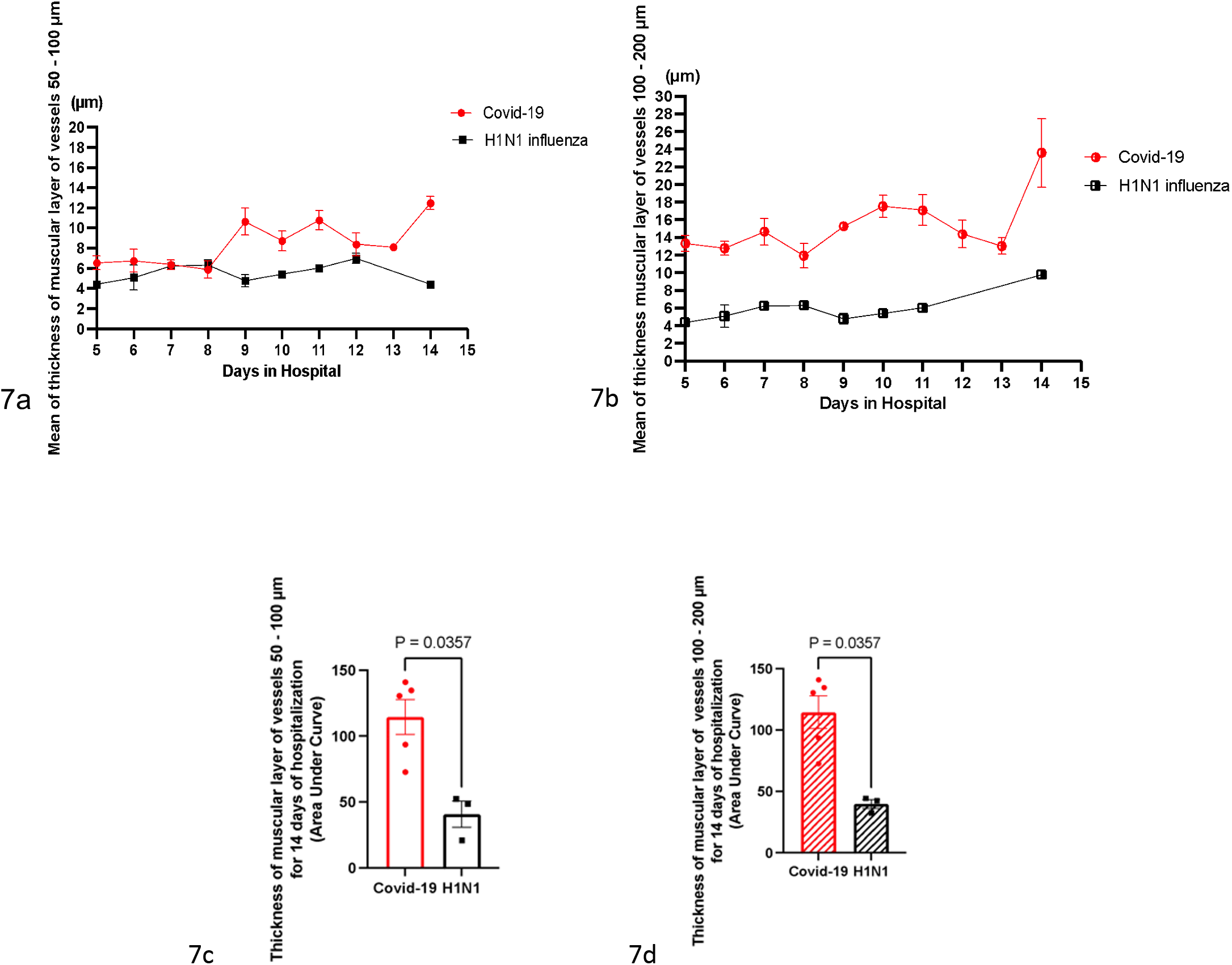
Graphs indicating correlation between the thickness of the pulmonary arterial muscular layer and the days in hospital in COVID-19 and H1N1 influenza. a - correlation between the thickness of the small vessels muscular layer and the days in hospital in COVID-19 and H1N1 influenza. b - correlation between the thickness of the medium-sized vessels muscular layer and the days in hospital in COVID-19 and H1N1 influenza. c - comparison of thickness of muscular layer of vessels 50 - 100 μm for 14 days of hospitalization (area under curve) between COVID-19 and H1N1 influenza (p=0.0357). d - comparison of thickness of muscular layer of vessels 100 - 200 μm for 14 days of hospitalization (area under curve) between COVID-19 and H1N1 influenza (p=0.0357).

When analyzing the dynamics of remodeling of the pulmonary arteries walls, a more pronounced thickening of the muscular layer was found in medium-sized vessels in COVID-19, compared with the H1N1 influenza, as well as a more rapid development of remodeling of medium-sized vessels rather than the small ones. A significant difference in the thickening of the muscular layer of small vessels between COVID-19 and H1N1 influenza was observed from the 8th day of hospitalization, while for medium-sized vessels this difference occurs in the earlier terms.

## Discussion

In the spring of 2020, we were stunned by the surge of infection by SARS-CoV-2 that resulted in a worldwide pandemic that influenced people in many ways. In the summer of 2020, we completed our initial study examining lung histology images from autopsy samples that were available in Ukraine. In that earlier study, we analyzed postmortem lung tissues collected from 10 patients who died of COVID-19 during the period of March to July 2020 and compared with patients who died by H1N1 influenza. While both viruses killed patients due to acute respiratory distress syndrome (ARDS), only COVID-19 patients exhibited thickened pulmonary vascular walls [13]. These findings promoted us to warn that COVID-19 patients may become predisposed to pulmonary arterial hypertension [15]. In this study, we extended these observations by using a larger number of patient samples to analyze this issue on pulmonary vascular remodeling in COVID-19.

By using 56 samples from patients who died by COVID-19, we confirmed our earlier study by showing that COVID-19 patients exhibit increased pulmonary vascular wall thickness and reduced lumen size, indicating the occurrence of pulmonary vascular remodeling and possibly pulmonary hypertension. The development of pulmonary vascular thickening took at least 14 days to develop after the onset of the disease. The expression of the SARS-CoV-2 spike protein was detected in both endothelial and smooth muscle cells of the pulmonary arteries of COVID-19 patients. Since the spike protein may interfere with the angiotensin regulation, angiotensin II-mediated growth of pulmonary vascular cells may play a mechanistic role. In addition, the spike protein may directly elicit cell growth signaling [13].

Our histopathological evaluations further provided mechanistic insights into the development of pulmonary vascular remodeling by showing; (i) the hypertrophy of the smooth muscle layer of the pulmonary arteries walls; (ii) perivascular fibrosis; (iii) edema and lymphostasis in the pulmonary arteries walls; (iv) inflammatory infiltration in the perivascular areas and in the pulmonary arteries walls; (v) perivascular hemosiderosis; (vi) neoangiogenesis in the perivascular areas; and (vii) vasa vasorum thrombosis.

A number of observational and case studies identified transthoracic echocardiogram- detectable pulmonary hypertension. As early as in the summer of 2020, Pagnesi et al. [19] published results of an observational study in Italy, concluding that the prevalence of pulmonary hypertension and right ventricular dysfunction in 200 COVID-19 patients was 12.0% and 14.5%, respectively. A Romanian study of 91 COVID-19 patients concluded the prevalence of pulmonary hypertension to be 7.69% and right ventricular dysfunction to be 10.28% [20]. Norderfeldt et al. [21] reported that 26 of 67 (39%) of COVID-19 patients exhibited acute pulmonary hypertension in Sweden. A 24-month follow-up study by the same group determined that the overall mortality was 61.5% in patients with acute pulmonary hypertension [22]. Numerous case studies also that reported the occurrence of pulmonary hypertension after contracting COVID-19 in various countries, including China [23], Netherlands [24], the United States [25], Pakistan [26], and Mexico [27]. The onset of pulmonary hypertension post COVID- 19 is unusually rapid, which is similar to two cases of COVID-19 vaccine-associated pulmonary hypertension [28], strengthening the possibility of the role of the spike protein in the pathogenic mechanism.

In summary, the present study presents a histopathological evaluation of the pulmonary vessels in patients who died of COVID-19, confirming that pulmonary vascular thickening and lumen narrowing occur in response to a SARS-CoV-2 infection. Since H1N1 patients who also died of ARDS do not exhibit pulmonary vascular remodeling, some unique features of SARS- CoV-2 specifically affect the pulmonary vessels as well as possibly vessels in general. Given that the spike protein is known to affect ACE2 that regulates angiotensin II levels, it is likely that this viral membrane fusion protein plays a critical role in the pathogenesis of the complications associated with COVID-19.

